# Mutational Crosstalk between Nuclear Driver Genes and Mitochondrial DNA in Gingivobuccal Oral Cancer

**DOI:** 10.1101/714519

**Authors:** Arindam Palodhi, Soumi Sarkar, Arindam Maitra

## Abstract

Somatic alterations in nuclear and mitochondrial genomes are known to play important role in the initiation and progress of tumours. However, crosstalk between these two genomic components in tumours have not been well investigated till date. We have excavated evidence of such crosstalk in gingivobuccal oral cancer. We have identified significant correlation between somatic mutations in nuclear driver genes and mitochondrial genes in 89 gingivobuccal oral cancer patients. Strong positive correlation between nuclear driver genes *CASP8, NOTCH1, USP9X, TRPM3* and mitochondrial genes has been observed. Patients having mutations in nuclear driver genes except *TP53*, were also shown to harbour higher mitochondrial mutation load. *TP53* mutations were shown to be negatively correlated with somatic mutations in mitochondrial *12S rRNA* and *ATP6*. Also, *TP53* mutations were negatively correlated with mutations in *NOTCH1, HRAS* and *USP9X*. We have identified a number of nuclear genes coexpressed with mitochondrial genes, although their association with tumorigenesis could not be established. Our observations suggest that in the absence of mutations in *TP53*, mutations in mtDNA and other driver genes might aid in the progression of cancer in a complementary fashion.

## Introduction

Eukaryotic cells meet their energy requirement by producing most of their ATP via the electron transport chain. This requires a high level of synchronization of the two obligate genomes, the mitochondrial genome (mtDNA) and the nuclear genome (nDNA) (Wolff et al., 2014). In addition to coding for the majority of the components of the electron transport chain (ETC), the nDNA also codes for all proteins required for the replication and upkeep of mitochondria. Therefore, mitochondrial-nuclear (mitonuclear) interaction at the genome level is essential for the assembly and function of the mitochondrial ETC (Ryan et al., 2007).

A large number of mitochondrial functions, including important metabolic activities are regulated by p53 (Lago et al., 2011). p53 mediated base-excision repair pathway protects mtDNA from oxidative damages caused by reactive oxygen species (ROS) (Kazak et al., 2012). p53 has also been found to relocate inside mitochondria and mediate apoptosis via mitochondrial membrane depolarization and caspase activation (Marchenko et al., 2000; Vaseva et al., 2012). These evidences indicate that the p53 might be involved in mitochondrial homoeostasis.

Oral squamous cell carcinoma of the gingivobuccal region (OSCC–GB) is the most prevalent cancer among males and fifth most prevalent among females from the Indian subcontinent (NICPR 2019). Of about 350,000 new cases that arise worldwide each year; 119,992 new cases are contributed by India in 2018 (Globocan 2018). Tobacco chewing in various forms, such as *betel quid, khaini* and *gutkha*, is traditionally popular in India and is known to enhance the risk of oral cancer (Gupta et al., 2011). Due to this particular mode of tobacco consumption, oral cancer lesions in the Indian patients are mostly localized in the gingivobuccal region, comprising buccal mucosa, retro-molar trigone and lower gum.

We have previously identified the nuclear driver genes (ICGC-India Project Team, 2013), important structural variations in chromosomes (Biswas et al., 2019) and potentially druggable pathways (Biswas et al., 2014) in OSCC-GB. We have also reported somatic driver mutations in mtDNA harboured by oral cancer patients (Palodhi et al., 2018). In this report, we provide information which might be useful in deciphering the interplay between the nDNA and the mtDNA in tumourigenesis. In order to understand this, we have studied the correlation between somatic mtDNA mutations and somatic mutations in the nuclear driver genes from 89 gingivobuccal oral cancer patients.

## Materials and Methods

Patients of OSCC-GB were recruited with informed consent (ICGC India Project Team, 2013). Tumour and blood samples were collected at the time of surgical excision, from treatment-naive OSCC-GB patients. Only patients with concordant histologic diagnosis by two independent reviewers were included. Sections of tumours that contained at least 80% tumour nuclei among total cellular nuclei were used. Mitochondrial and nuclear DNA mutations were identified from paired DNA samples from tumour and matched blood from 89 patients. Whole genome sequencing was performed for 25 patients and mitochondrial DNA sequences were extracted from the same. For the next 64 patients, mitochondrial genome was amplified using long range amplification and sequenced in-house (**Supplementary Table S1**). Mutations in the nuclear genome were identified from whole genome and whole exome sequencing data of 25 and 64 patients respectively (**Supplementary Table S2**). Sample BAM files used in this study are deposited in the European Genome Phenome Archive under accession code EGAS00001001901 (WGS), EGAS00001000249 & EGAS00001001028 (WES), EGAD00001004987 (mtDNA sequence) and EGAS00001003285 (RNA-seq).

The degree of correlation between mutation events in nuclear driver genes and mtDNA was assessed by calculating Phi-coefficient. Spearman’s test was used to assess the correlation between mitochondrial mutation load and nuclear driver mutation. All statistical analyses were performed using R (R-core team, 2014).

## Results

Ten nuclear genes (*TP53, FAT1, CASP8, USP9X, MLL4, NOTCH1, HRAS, UNC13C, ARID2* and *TRPM3*) frequently mutated in OSCC-GB have been reported previously (ICGC India Project Team, 2013). Among the 89 patients, 55 patients (61%) harboured somatic mutations in *TP53. FAT1* or *CASP8* were mutated in 26 patients each, followed by *NOTCH1* (19 patients). Mutations in the mitochondrial genome in these patients were categorized into 17 features which are as follows; 13 mitochondrial coding genes as 13 individual features, 2 rRNAs (12S and 16S) and the *D-loop* region each as one feature and all mitochondrial tRNAs as a single feature. We calculated the Phi-coefficient between the somatic mutations in these nuclear driver genes and those in the mtDNA (**Supplementary Table S3**). We found that mutations in *ARID2* positively correlated with mutations in *D-loop, mt-tRNA* and *ND4*. Somatic mutations in four nuclear driver genes, *CASP8, NOTCH1, USP9X and TRPM3* displayed positive correlation with mutations in mitochondrial genes *COX3, ND5, ND1* and *COX2* respectively. Mutations in *TP53* were found to be negatively correlated with somatic mutations in mitochondrial 12S rRNA and *COX3* gene. Additionally we observed that *TP53* mutations were negatively correlated with mutations in *NOTCH1, HRAS* and *USP9X*. Thus *NOTCH1* and *USP9X* simultaneously appeared to be negatively correlated with *TP53* mutations and positively correlated with some mitochondrial gene mutations **(Figure 1A)**. Mutations in rest of the driver genes, namely *FAT1, HRAS, UNC13C* and *MLL4* were not significantly correlated with somatic mtDNA mutations.

**Figure 1:**
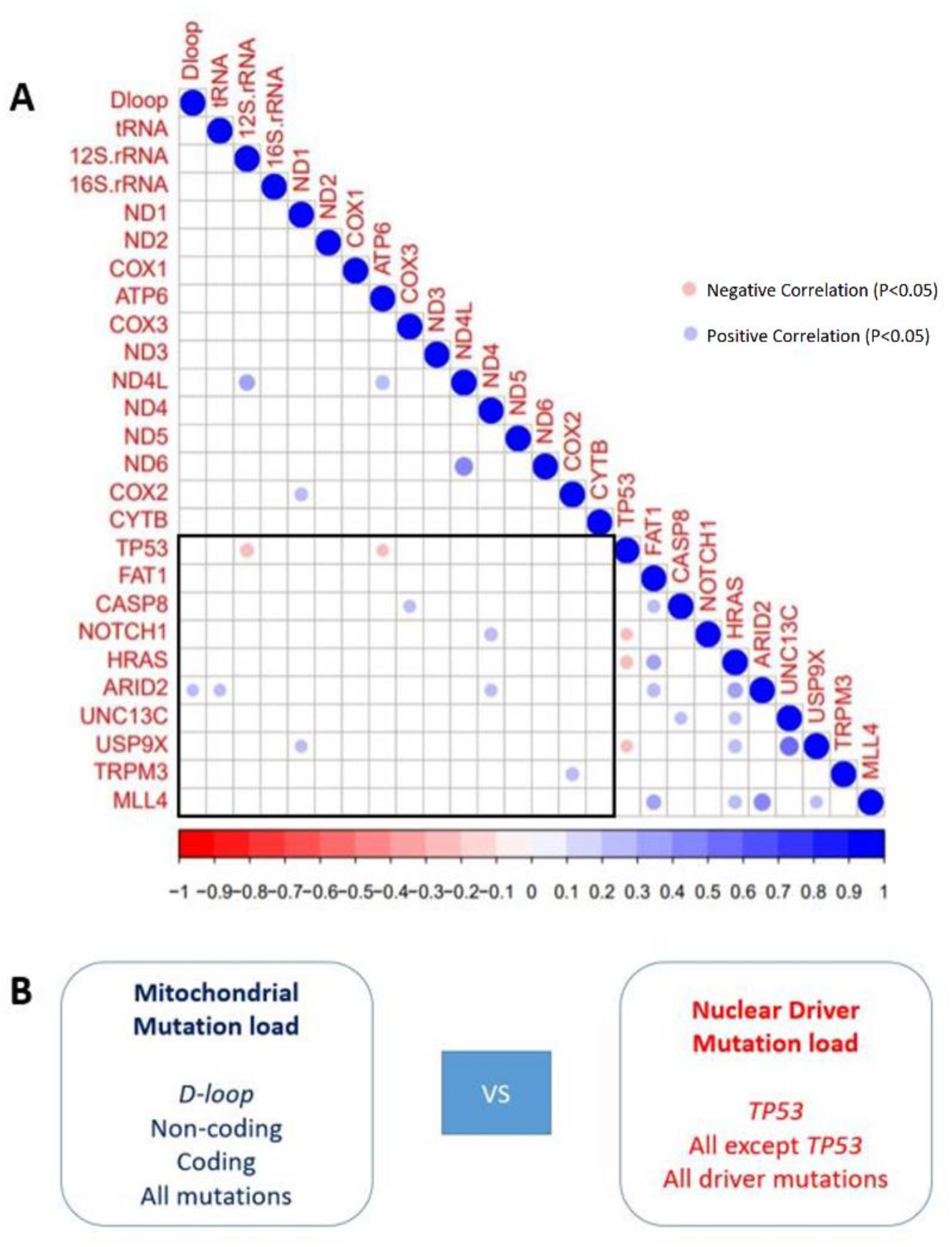
(A) Correlation between mitochondrial and nuclear mutations: Red circles indicate negative correlation and blue circles indicate positive correlation. Only significantly correlated genes are reported. (B) Schematic diagram representing the correlated mutation burden groups: Mutation loads in mitochondrial *D-loop* region, non-coding genes, coding genes and all of these mutations combined were compared with mutations load in nuclear driver genes, *TP53* gene and all genes except *TP53* to analyse mito-nuclear interaction at genome level.

We also investigated the relationship between the mutation load of nuclear driver genes and mutation load of mtDNA in our dataset. Some of the nuclear driver genes, most notably *TP53* has tumour suppressor activity. Loss of function mutations in *TP53* may lead to higher somatic mutation load in mtDNA, making the guardian of the mitochondrial genome (Park et al., 2016). We have categorized mutation burden of nuclear driver genes in each patient in 3 groups; mutation burden in all driver genes, mutation burden in *TP53* and mutation burden in 9 other non-*TP53* driver genes. Similarly, total somatic mutation burden in mtDNA was categorized into 4 groups; cumulative mutation burden in mitochondrial protein coding genes, mutation burden in *D-loop* as this is the control region for transcription and translation of mtDNA, mutation burden in all non-coding regions which includes *D-loop*, 2 mitochondrial rRNAs and 22 mitochondrial tRNAs, and total mutation burden which combines all the categories mentioned **(Figure 1B)**. Spearman correlation test was performed between each of the nuclear mutation groups and individual mtDNA mutation groups (**Supplementary Table S4**). The mtDNA coding mutation load and mutation burden in 9 non *TP53* driver genes was found to be significantly positively correlated (ρ = 0.2700105 P = 0.02087).

## Discussion

We have observed that patients harbouring mutations in other nuclear driver genes such as *HRAS, NOTCH1, USP9X* etc., are more likely to harbour mutations in mitochondrial genes. We also found that most of the patients carrying mutations in *TP53* are less likely to harbour mutations in mtDNA. We hypothesize that in the absence of mutations in *TP53*, mutations in mtDNA and other driver genes might aid in the progression of cancer in a complementary fashion. The positive correlation between the mutation load in mtDNA and mutation burden in non-*TP53* driver genes further supports our hypothesis.

In this study, we have also investigated the co-expression of 1701 nuclear coded genes (gene list in **Supplementary Table S5**) which are associated with structure and function of mitochondria and mtDNA coded respiratory complex genes (result not shown). This analysis did not throw any light on the retrograde signalling pathway which might be associated with oral cancer. This phenomenon might be attributed to factors such as post-transcriptional regulation of mitochondrial RNA processing, which result in the difference in abundance of 13 mitochondrial transcripts (Rorbach et al., 2012; Palodhi et al., 2018), despite that they are transcribed at once as a single poly-cistronic RNA (Mercer et al., 2011). Previous studies have also indicated additional roles played by translational modulation of nuclear and mitochondrial genes in regulating the mitochondrial and nuclear OXPHOS protein levels which might be necessary for the various subunits of the electron transport chain to assemble at a specific stoichiometric ratio (Couvillion et al., 2016). A recent report by Bajzikova et al. (2018) suggest that the presence of functional mitochondria is necessary for the requirement of pyrimidine biosynthesis in tumour cells. The above reports along with our observations point towards the possible shortcoming of gene expression data as surrogate for mito-nuclear interaction at RNA level. Although altered expression of mitochondrial and nuclear OXPHOS genes independently, might indicate potential reprograming of the tumour metabolism (Reznik et al., 2017). In some cases, the correlation between the nuclear and mitochondrial gene expression may have yet unknown implications for the respiratory activity.

Here we report positive correlation between mtDNA mutations and mutation burden in non-*TP53* driver genes in gingivobuccal oral cancer. We also observed negative correlation between *TP53* mutation and mutations in the mtDNA in cancer which has not been reported earlier. These observations suggest that in the absence of mutations in *TP53*, mutations in mtDNA and other driver genes might aid in the progression of cancer in a complementary fashion. The possible mechanism via which mtDNA coding mutations and mutations in 9 non-*TP53* driver genes aid in tumour progression in absence of *TP53* loss of function mutations needs further investigation.

## Acknowledgements

We acknowledge the ICGC-India Project team and the Core Technologies Research Initiative (CoTeRI) of NIBMG for generation of the sequencing data. We thank Dr. Sillarine Kurkalang for her help in reviewing this manuscript. We are grateful to the Department of Biotechnology, Ministry of Science and Technology, Government of India, for providing financial and logistical support to the ICGC – India Project.

